# Crystal structure of activated MAPKAPK2 kinase reveals a C-lobe allosteric pocket for furan-2-carboxamide inhibitors

**DOI:** 10.64898/2026.10.01.755332

**Authors:** Edward Beaumont, Stephanie Duclos, Sandrine Thieffine, Rahul Yadav, John Barker

## Abstract

MAPK-activated protein kinase 2 (MAPKAPK2, MK2) is a downstream substrate of p38 MAPK that regulates inflammatory signaling and cytokine synthesis, making it an attractive therapeutic target. Although several crystal structures of MAPKAPK2 with ATP-competitive inhibitors have supported drug discovery, structural elucidation of selective non-ATP-competitive inhibitors has remained challenging. MK-25 is a non-ATP-competitive furan-2-carboxamide MAPKAPK2 inhibitor developed at Merck to overcome the selectivity and cellular-efficacy limitations of ATP-competitive chemotypes. Here, we show through isothermal titration calorimetry that MK-25 binds the activated MAPKAPK2, consistent with state-dependent allosteric binding. To determine the binding mode of MK-25, we solved a 2.8 Å resolution crystal structure of an activated MAPKAPK2 construct in ternary complex with the ATP-site inhibitor PF-3644022 and a bromo-analogue of MK-25. The ligand occupies a pocket in the C-lobe overlapping the protein-substrate recognition region and formed through rearrangements of αD and αG helices and the activation segment. These interactions are consistent with the structure-activity relationships reported for the furan-2-carboxamide chemotype and identify a ligandable allosteric pocket that may facilitate future structure-guided design of selective, non-ATP-competitive MAPKAPK2 inhibitors.

## 1. Introduction

The p38 mitogen-activated protein kinase (MAPK) pathway controls cellular responses to stress and pro-inflammatory stimuli through phosphorylation of downstream substrate proteins (Kyriakis et al., 2001; Cuenda et al., 2007). Among these, MAPKAPK2 has attracted particular interest as a target in autoimmune and inflammatory diseases because it regulates the production of inflammatory mediators including TNFα and interleukins indirectly controlling gene expression at the level of translation by phosphorylation of RNA-binding proteins. (Menon et al., 2018; Beamer et al., 2021).

Direct inhibition of p38 MAPK has been pursued extensively, but clinical development has been limited by compound toxicity and insufficient efficacy (Schindler et al., 2007; Fiore et al., 2016). These limitations have motivated efforts to target MAPKAPK2, with the aim of improving selectivity and safety while modulating the p38 MAPK pathway. However, the development of MAPKAPK2 inhibitors has proven challenging. ATP-competitive chemotypes can achieve potent biochemical inhibition yet often suffer from limited kinase selectivity and reduced cellular efficiency because they must compete with high intracellular ATP concentrations (Fiore et al., 2016). Accordingly, non-ATP-competitive inhibition represents an attractive alternative strategy for obtaining selectivity and improved functional activity.

MAPKAPK2 belongs to the Ca^2+^/calmodulin-dependent protein kinase (CaMK) superfamily and is regulated by a distinctive activation mechanism where phosphorylation induces a relief of autoinhibition (Manning et al., 2002; Ben-Levy et al., 1995; Underwood et al., 2003). Human MAPKAPK2 contains an N-terminal region, a central kinase domain, and a C-terminal regulatory segment that includes an autoinhibitory helix (Aiα) in Figure 1A (Meng et al., 2002; Zu et al., 1995). Phosphorylation by p38α at Thr222, Ser272, and Thr334 promotes displacement of the autoinhibitory helix and stabilizes the active kinase conformation (Ben-Levy et al., 1995; Underwood et al., 2003; Fiore et al., 2016). Several crystal structures of MAPKAPK2 in inactive and ATP-competitive inhibitor-bound states have provided valuable insight into kinase regulation and orthosteric ligand recognition (Meng et al., 2002; Underwood et al., 2003; Anderson et al., 2009a and 2009b).

**Figure 1.**
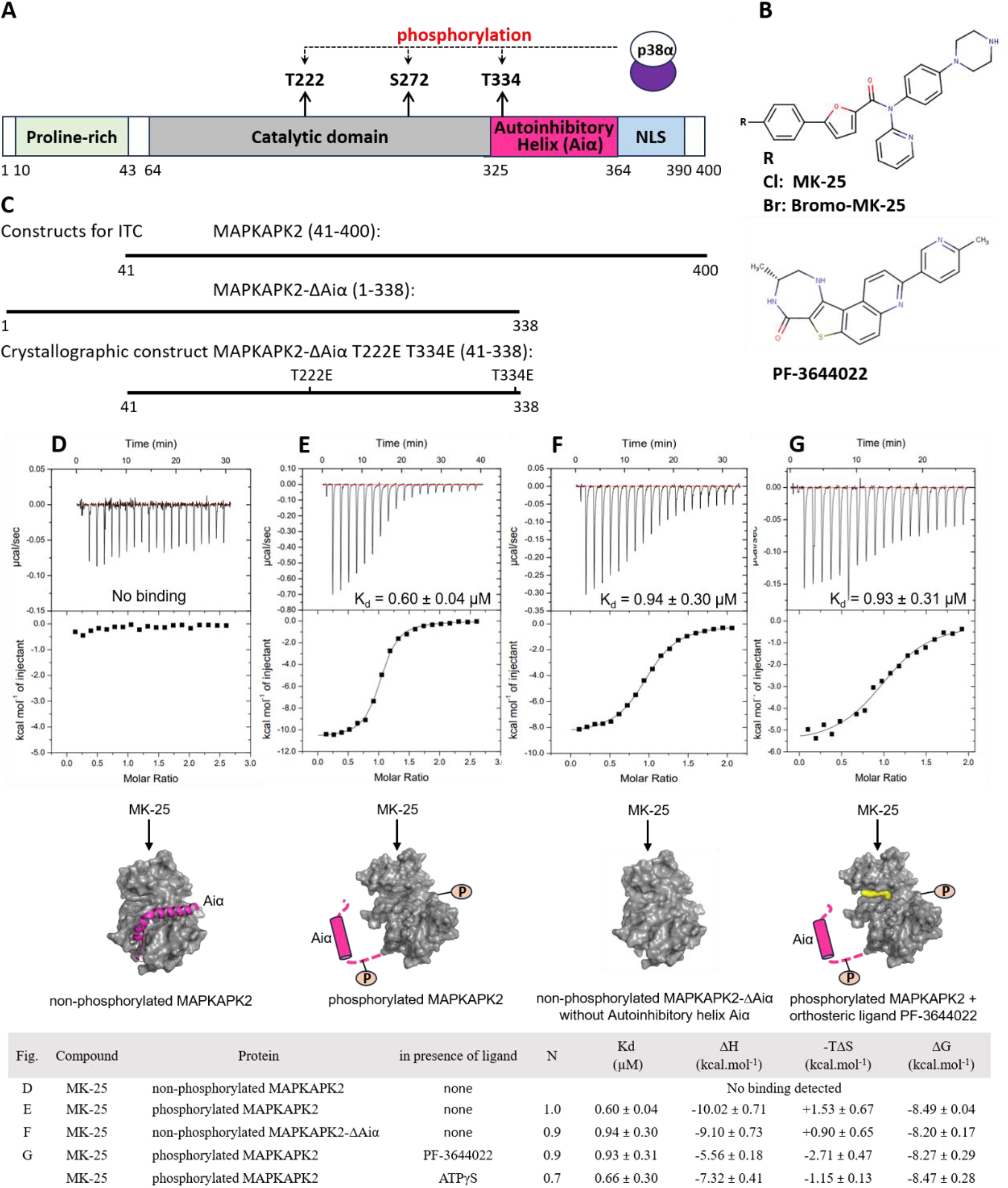
MK-25 binds specifically to phosphorylated MAPKAPK2 at high affinity. (A) Scheme of the domain organization of MAPKAPK2. Residue range for each domain is labelled under the domain diagram according to the Uniprot entry P49137. (B) Chemical structure of MK-25, Bromo-MK-25 and PF-3644022. (C) The boundaries of the MAPKAPK2 constructs used in this study, in relation to the schematic diagram in (A): MAPKAPK2 (Q41-H400) and MAPKAPK2-ΔAiα (M1-T338) and the crystallographic construct MAPKAPK2 (Q41-T338 T222E T334E). (D, E, F, G) Representative ITC thermograms for MK-25 to different MAPKAPK2 states. Autoinhibitory segment (Aiα) is represented in pink, and the orthosteric ligand is shown in yellow (D) Titration of MK-25 into non-phosphorylated MAPKAPK2. (E) Titration of MK-25 into phosphorylated MAPKAPK2. (F) Titration of MK-25 into non-phosphorylated MAPKAPK2-ΔAiα. At least two titrations were performed for each experiment, and the averages and errors are reported. (G) Titration of MK-25 into phosphorylated MAPKAPK2 in presence of orthosteric ligand PF-3644022.

A notable advance in the development of allosteric inhibitors was the affinity-based discovery of a diaryl furan-2-carboxamide series that was further optimized to yield compound 25 (MK-25) (Figure 1B). MK-25 was found to be a selective non-ATP-competitive MAPKAPK2 inhibitor (Huang et al., 2011). Biochemical and NMR analyses supported binding away from the ATP site, and exploration of the structure-activity relationships (SAR) across the scaffold, highlighting the substituted furan-2-carboxamide core and the distal aryl-piperazine substituents as critical moieties for potency and activity (Huang et al., 2011; Huang et al., 2012). Despite these advances, the precise binding site of this chemotype to the activated state of MAPKAPK2 has not been described.

Here, we combined isothermal titration calorimetry (ITC) and X-ray crystallography to define the interaction of MK-25 analogues with MAPKAPK2. ITC shows that MK-25 binds to the activated form of MAPKAPK2, consistent with the activated state-dependent recognition. We then solved the crystal structure of a constitutively active MAPKAPK2 kinase-domain construct, two phosphomimetic mutations T222E T334E in ternary complex with the ATP-competitive inhibitor PF-3644022 (Anderson et al., 2009b; Mourey et al., 2010) and a bromo-analogue of MK-25. The structure reveals an allosteric pocket in the MAPKAPK2 C-lobe, overlapping the protein substrate-recognition region, that is formed through conformational rearrangements upon kinase activation. These data provide a structural explanation for the SAR reported across the furan-2-carboxamide series (Huang et al., 2011; Huang et al., 2012) and offers a foundation for structure-guided design of selective non-ATP-competitive MAPKAPK2 inhibitors.

## 2. Materials and Methods

### 2.1. General information

PF-3644022 and ATPγS were purchased from Sigma-Aldrich. MK-25 and Bromo-MK-25 were purchased from Cayman Chemical and synthesized based on Huang et al., 2011, respectively.

### 2.2. Protein constructs

The synthetic genes of the recombinant human MAPKAPK2, and upstream kinases MKK6-DD and murine p38α, with a TEV-protease cleavable 6xHis tag fused to their N-terminus, were cloned as BamH1-Xho1 fragment into pTRiJ-HV vector (Genscript). Genes MAPKAPK2 Q41-T338 T222E T334E, MAPKAPK2 H47-H400 and of wild type p38α were codon-optimized for *E*.*coli* expression, and genes of truncated MAPKAPK2 M1-T338 and of MKK6-DD were codon optimized for insect expression (protein sequences in supplementary information).

### 2.3. Protein expression

MAPKAPK2 (H47-H400 and Q41-T338 T222E T334E) were expressed in BL21(DE3) and p38α in BL21(DE3) pLysS in TB media (Teknova) and grown at 37 °C until an OD_600_ of 0.7-0.8 then induced with 0.4 mM IPTG and left to express overnight at 20 °C. Cells were harvested at 4000g for 15 min. MAPKAPK2 (M1-T338) and MKK6-DD were expressed in Sf21 cells using *flash*BAC ULTRA expression system (Oxford expression technologies). Cells were cultured at 27°C to a density of 1.4 x 10^6^ cells ml^-1^ and infected at an MOI of 1 in 10 L of SF900-II media (Gibco) in wave bag. At 68 hrs post-infection, cells were harvested at 1600g for 15 min.

### 2.4. Protein purification

For variants of MAPKAPK2 (H47-H400, M1-T338 and Q41-T338 T222E T334E) and MKK6-DD; cells were suspended in lysis buffer containing 50 mM Tris-HCl pH 7.5, 300 mM NaCl, 1 mM DTT, 20 mM imidazole, 5% glycerol, supplemented with benzonase (Sigma-Aldrich) and protease inhibitor tablets (Roche). Cell disruption was performed by sonication, and the lysate was clarified by centrifugation at 15000 rpm and 4 °C for 1 hr. The supernatant was loaded onto a HisTrap FF crude column (Cytiva). The protein was gradually eluted with an imidazole gradient using elution buffer (50 mM Tris-HCl pH7.5, 300 mM NaCl, 1 mM DTT, 500 mM imidazole). The protein was dialysed in the presence of 6His-TEV protease (1:100 w/w) in buffer (50 mM Tris-HCl pH 7.5, 300 mM NaCl, 1 mM DTT and 5% glycerol) at 4°C, overnight. The sample was further purified via subtractive IMAC using a HisTrap HP column (Cytiva), followed by gel filtration (Superdex 75 26/60 column, (Cytiva) with 20 mM HEPES pH7.5, 100 mM NaCl, 5 mM MgCl_2_, 1 mM TCEP as running buffer before concentration to 8.9 mg ml^-1^ (MAPKAPK2 Q41-T338 T222E T334E), 8.5-9.5 mg ml^-1^ (MAPKAPK2 H47-H400 and M1-T338) and 1 mg ml^-1^ (MKK6-DD).

p38α purification was performed according to White et al. 2007.

### 2.5. Phosphorylation of p38α and MAPKAPK2

Constitutively active MKK6-DD was used to phosphorylate p38α. Then, phosphorylated p38α was then used to phosphorylate MAPKAPK2.

The phosphorylation of p38α and MAPKAPK2 was performed using a common protocol. Each kinase and its upstream activator were dialyzed separately in phosphorylation buffer (25 mM HEPES pH 7.0, 100 mM NaCl, 20 mM MgCl_2_, 2 mM DTT, 0.5 mM EDTA, 10% glycerol) for 2 h at 4 °C, then mixed, diluted to 30 ml, and incubated with 10 µM sodium orthovanadate and 4 mM ATP (pH 7.0) for 2 h at 22°C. Phosphorylation state was monitored by mass spectrometry.

For p38α activation, 10 mg of p38α (4 mg ml^-1^) was incubated with 1 mg of MKK6-DD (1 mg ml^-1^). The reaction was purified on a Resource Q column (Cytiva) using a wash buffer (20 mM Tris pH 8.0, 2 mM TCEP) and elution buffer (20 mM Tris pH 8.0, 500 mM NaCl, 2 mM TCEP), then dialysed in buffer (25 mM HEPES pH 7.5, 50 mM KCl, 2 mM TCEP, 5% glycerol) and concentrated to 2 mg ml^-1^.

For MAPKAPK2 activation, 10 mg of MAPKAPK2 (4 mg ml^-1^) was incubated with 2 mg of the phosphorylated p38α (2 mg ml^-1^). The reaction was purified on a MonoS 5/50 column (Cytiva) using a wash buffer (50 mM HEPES pH 7.5, 100 mM NaCl, 2 mM TCEP, 10% glycerol) and elution buffer (50 mM HEPES pH 7.5, 1 M NaCl, 2 mM TCEP, 10% glycerol), then dialysed in buffer (25 mM HEPES pH 7.5, 50 mM KCl, 2 mM TCEP, 5% glycerol) and concentrated to 2 mg ml^-1^.

### 2.6. Isothermal titration calorimetry

MAPKAPK2 was dialysed at 4°C over night against 20 mM HEPES pH 7.5, 100 mM NaCl, 5 mM MgCl_2_ and 1 mM TCEP. The dialysis buffer was then degassed at room temperature. Titrations were carried out at 25°C, using an ITC200 calorimeter (MicroCal Inc). The small molecule (20 to 30 µM) was loaded into the sample cell and the small molecule was loaded into the syringe (200 to 300 µM in degassed dialysis buffer with 2% DMSO). For experiments performed in the presence of PF-3644022, the compound was included in both the sample cell and syringe at a final concentration of 50 µM. The data were then analysed using Origin (OriginLab Corp.) and fitted to a single-site binding model implemented in the software of the instrument supplier.

### 2.7. Crystallization and X-ray data collection

MAPKAPK2 co-crystals were grown by hanging drop vapour diffusion at 22°C by mixing equal volumes of protein (8.9 mg/ml) with precipitant (1.3-1.5 M NaMalonate, 0.1 M HEPES pH6-8, 0.5% Jeffamine ED2003). Both PF-3644022 and Bromo-MK25, at 1 mM final concentration, were incubated together with MAPKAPK2 at room temperature for 1 hour. Prior to freezing in liquid nitrogen, crystals were cryoprotected by brief transfer to a solution of reservoir solution supplemented with 25% (v/v) glycerol. Diffraction data were collected on the I03 beamline at Diamond Light Source (Oxford, U.K.). A complete dataset was collected from a single crystal under a cryogenic stream at 100 K.

### 2.7. Structure determination

The diffraction dataset was processed using xia2 DIALS pipeline (Winter et al., 2018), which uses XDS (Kabsch et al., 2010), Pointless (Evans, 2006) and Aimless (Evans and Murshudov, 2013) of the CCP4 suite (Winn et al., 2011). The structure of MAPKAPK2 complex was solved by molecular replacement using the previously solved structure (PDB 1NY3, Underwood et al., 2003) as a reference model. The refinement was performed with BUSTER (Bricogne et al., 2017) and manually corrected using COOT (Emsley et al., 2010). Compound dictionaries were generated using AFITT (Wlodek et al., 2006). The diffraction data and refinement statistics are shown in Table **T1**.

## 3. Results

### 3.1. MK-25 binds selectively to activated MAPKAPK2

To define the conformational requirements for the binding of MK-25, we first measured the interaction of MK-25 with several MAPKAPK2 constructs by ITC (Figure 1C-G). The constructs comprised the MAPKAPK2 kinase-domain construct (residues H47–H400), which retains the C-terminal autoinhibitory helix (Aiα), and the truncated MAPKAPK2-ΔAiα (residues M1–T338), in which Aiα is removed; the first was studied in non-phosphorylated and p38α-phosphorylated forms (phosphorylation likely at Thr222, Ser272, and Thr334), allowing the contributions of autoinhibition and phosphorylation-dependent activation to be assessed independently.

No detectable binding was observed for the non-phosphorylated kinase construct, indicating that the autoinhibited state does not support MK-25 binding, consistent with the tethered autoinhibitory helix (Aiα) occluding the binding site (figure 1D). In contrast, binding was observed with phosphorylated, p38α-activated MAPKAPK2 with submicromolar affinity (Kd = 0.60 ± 0.04 µM), indicating that MK-25 selectively binds the activated form of MAPKAPK2 (figure 1E).

To test whether displacement of the C-terminal autoinhibitory region is sufficient to expose the binding site, we next examined a constitutively active MAPKAPK2 construct lacking the C-terminal autoinhibitory helix Aiα (non-phosphorylated construct MAPKAPK2-ΔAiα, figure 1F). MK-25 also bound this construct with comparable affinity (Kd = 0.94 ± 0.30 µM). Together, these data support a model in which MK-25 recognizes a binding site that becomes accessible upon activation-associated rearrangement of the MAPKAPK2 C-lobe. Binding was also observed in the presence of both ATPγS and the ATP-competitive inhibitor PF-3644022 confirming a non-ATP-competitive mechanism (figure 1G).

### 3.2. Structure determination of an activated MAPKAPK2 ternary complex

To characterize the allosteric binding site and define the structural basis for ligand recognition, we determined the crystal structure of a constitutively active MAPKAPK2 kinase-domain without the autoinhibitory helix in complex with PF-3644022 and a bromo-substituted analogue of MK-25 (Bromo-MK-25) at 2.8 Å resolution (Figure 2A). The Bromo-substitution in MK-25 was used to allow anomalous diffraction to confirm the binding site if required. The crystallographic construct has the same N- and C-terminal boundaries as the protein used to obtain the initial furan-2-carboxamide hit (Huang et al., 2011); lacking the C-terminal autoinhibitory helix (Aiα), it carries phosphomimetic substitutions (T222E and T334E) to favor an active-like kinase conformation (Figure 1C).

**Figure 2.**
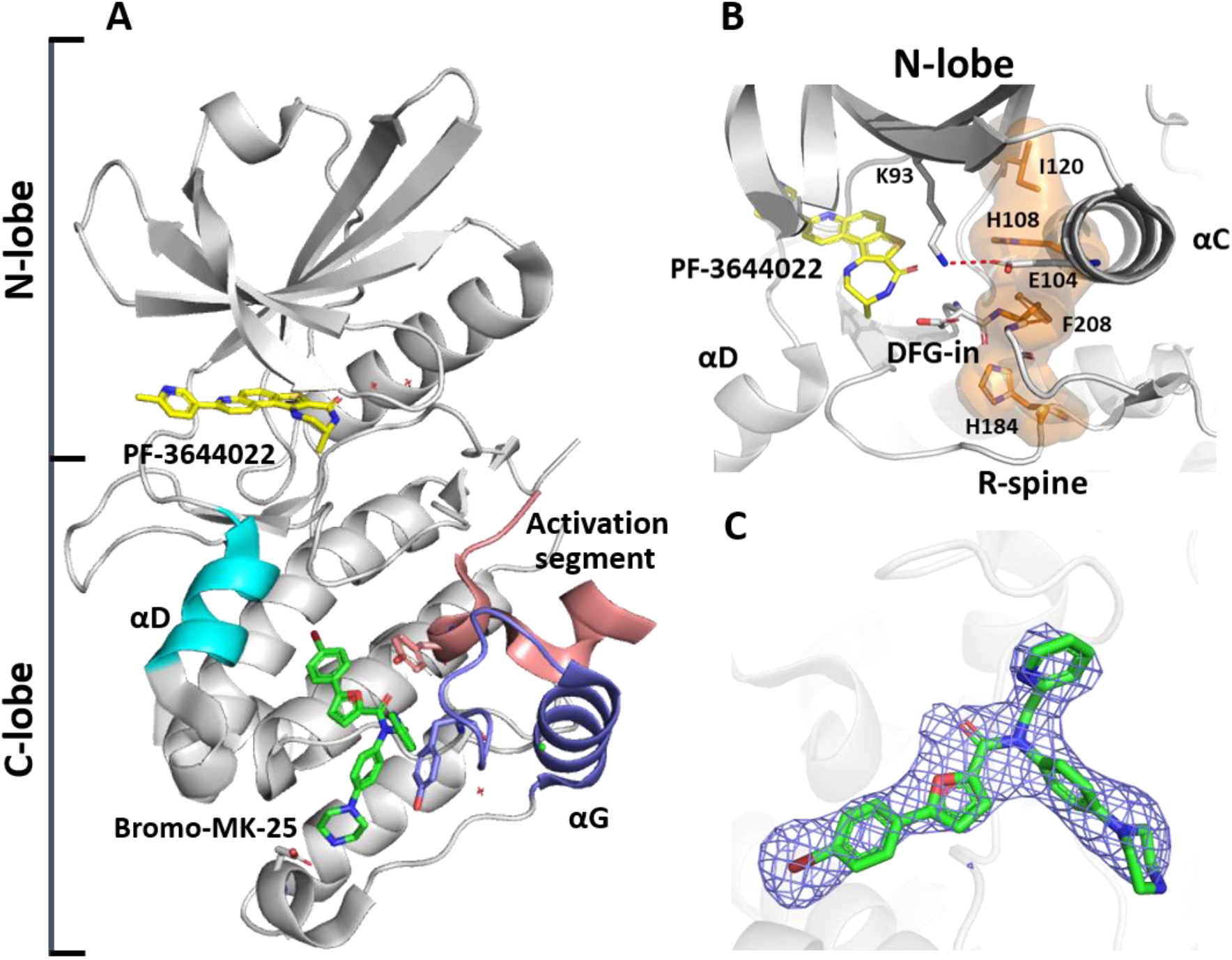
Crystal structure of the MAPKAPK2 complexed with PF-3644022 and Bromo-MK-25. (A) overall structure of activated MAPKAPK2 represented in grey cartoon. PF-3644022 and Bromo-MK-25 are shown in yellow and green sticks respectively. The secondary structures of interest are represented in cyan for αD, blue for αG and brown for the activation segment. (B) The N-lobe of MAPKAPK2 adopts an active conformation based on the orientation of key motifs, including the DFG-in, αC-in, R-spine in orange and the salt bridge Lys93-Glu104 interaction as a red dash line. (C) 2Fo-Fc electron density map in blue mesh is shown contoured at 1σ around Bromo-MK-25.

Crystals of the binary MAPKAPK2:Bromo-MK-25 complex failed to yield diffraction-quality data. The addition of the potent ATP-competitive inhibitor PF-3644022 (Anderson et al., 2009b) proved essential, enabling determination of the MAPKAPK2:PF-3644022:Bromo-MK-25 ternary complex structure at 2.8 Å resolution with R/R_free_ values of 22.7/24.8% (Table T1).

Unsurprisingly, the overall structure reveals an active-like kinase conformation. In the N-lobe, key hallmarks of an active protein kinase (Röhm et al., 2020; Taylor et al., 2011) are preserved, including the salt bridge formed by Lys93 and Glu104, αC-in positioning and DFG-in arrangement, together supporting assembly of the regulatory spine, composed of Ile120, His108, Phe208, His184 (Figure 2B). Electron density for the orthosteric inhibitor PF-3644022 is well defined in the ATP site, and its binding pose is similar to that reported in crystal MAPKAPK2 structure 3FYJ (Anderson et al., 2009b). Additional electron density in the C-lobe identifies the bound bromo-MK-25 analogue (Figure 2C). This ternary complex provides direct structural evidence that the furan-2-carboxamide chemotype binds outside the ATP pocket and instead occupies a distinct site in the C-lobe of activated MAPKAPK2.

### 3.3. The MK-25 analogue occupies a C-lobe pocket overlapping the substrate-recognition surface

A single molecule of the bromo-MK-25 analogue binds to MAPKAPK2, in a shallow but extended C-lobe pocket (Figure 3A and S1). The ligand lies across a largely hydrophobic surface formed by residues from αD, αG, and the activation segment (Figure 3A). The furan-2-carboxamide core adopts the *E*-conformer (cis-amide), consistent with reports that the *E*-conformer is favoured by 2-3 kcal/mol over the Z-conformer (Galan et al., 2013; Itai et al., 1992) (Figure 3B). The furan-2-carboxamide core is stabilized by two hydrogen bonds: one between the amide carbonyl and the hydroxyl of Tyr228, and another between the furan and the carbonyl of Gly259. These anchoring interactions are consistent with those previously proposed to contribute to the potency of this series (Huang et al., 2011).

**Figure 3.**
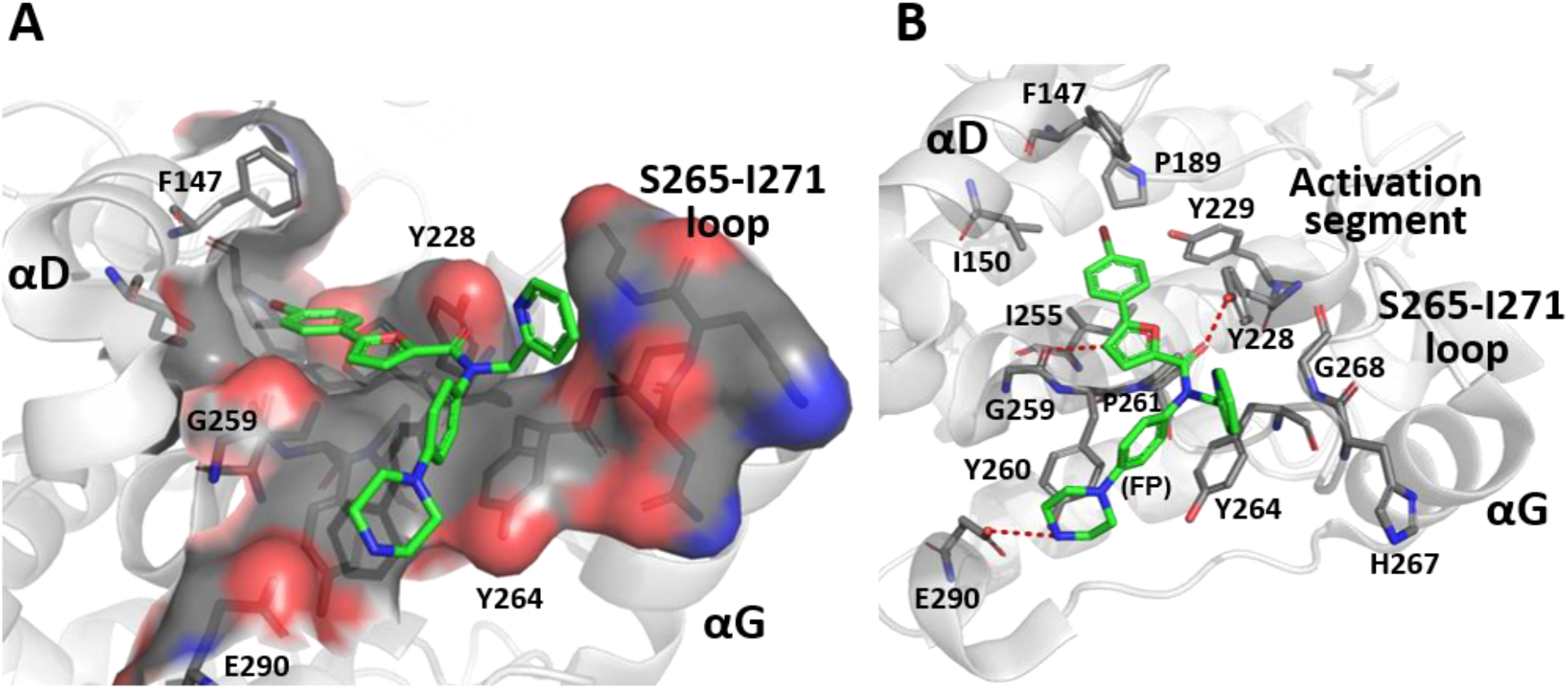
Binding of Bromo-MK-25 to MAPKAPK2. Bromo-MK-25 is represented in green sticks in all panels. (A) Surface representation of Bromo-MK-25 bound MAPKAPK2 binding site. Red, blue and grey colour patches correspond to oxygens, nitrogen and carbon, respectively. (B) The key binding site residues are shown and labelled. Hydrogen bonds between Bromo-MK-25 to the protein are shown as dashed lines.

The neighboring para-bromo-phenyl group occupies a shallow hydrophobic pocket formed by residues including Ile255, Ile150, Pro189, and Phe147. On the other branch of the main scaffold, the phenyl group (FP; Figure 3B) is twisted to avoid internal clash with the furan moiety and forms a tilted T-shape π-π interaction with Tyr264. The following para-piperazine interacts with Tyr260 via H-π interactions, helping to orient the terminal basic amino group of the piperazine toward Glu290 to form a salt bridge with the side chain carboxyl group. The 1-phenyl-piperazine substituent was associated with both binding affinity and cellular activity (Huang et al., 2012). Finally, the 2-pyridyl ring is positioned to form a π-stack with the amide backbone of His267-Gly268 from the Ser265-Ile271 loop. It is positioned consistent with its previously reported contribution to potency within this series (Huang et al., 2011). The observed binding mode is consistent with the SAR previously reported for the furan-2-carboxamide series (Huang et al., 2011; Huang et al., 2012).

### 3.4. Activation-associated remodeling of the MAPKAPK2 C-lobe creates the allosteric pocket

Comparison with known inactive MAPKAPK2 structures shows that the formation of the allosteric site requires substantial reorganization within the C-lobe (Figure 4). The most prominent differences involve the αD and αG helices, the activation segment, and the C-terminal region adjacent to the autoinhibitory helix. In the activated structure complex, residues 330-338 are disordered, consistent with release of the autoinhibitory helix and with the construct design used for crystallization (i.e. deletion of the subsequent autoinhibitory helix (Leu336-Asp366). Concomitantly, the activation segment folds into the newly accessible cradle of a neighbouring molecule of MAPKAPK2 (in *trans*), left vacant by the absence of the autoinhibitory helix, similar to what has been observed in several constitutively active MAPKAPK2 crystal structures (Underwood et al., 2003; Hillig et al., 2007). Moreover, the αD helix is rotated to partially occupy the volume left vacant by the absent autoinhibitory helix and to flank the αF helix. Finally, the αG helix and the adjacent loop Ser265-Ile271 become ordered, compared to the inactive conformation, and pack against the side of the displaced activation segment. These rearrangements are compatible with regulatory behavior described for related CaMK superfamily kinases, including the αD rearrangement and activation-associated displacement of the autoinhibitory segment observed in CaMKII (Rellos et al., 2010).

**Figure 4.**
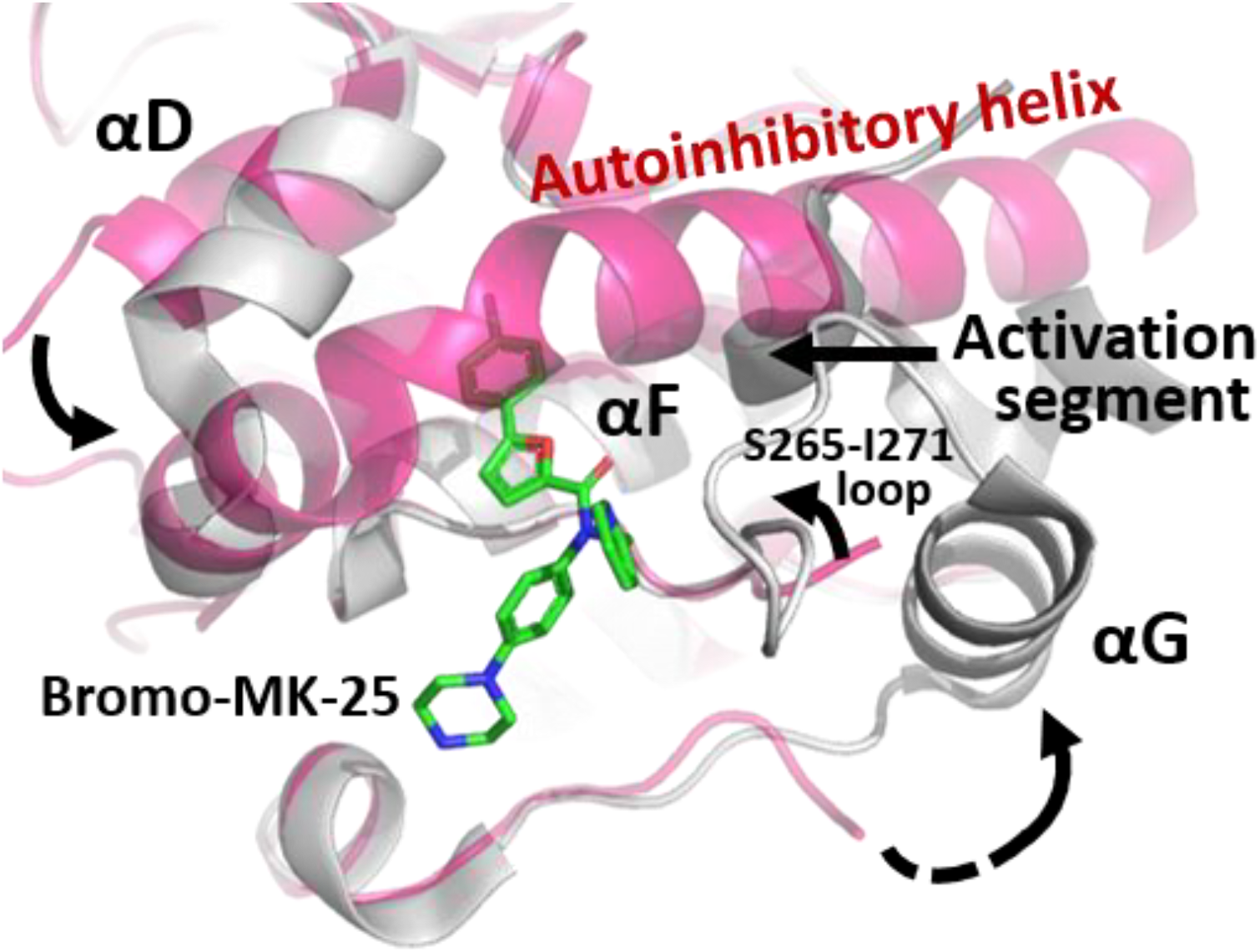
Conformational changes that create the allosteric pocket. Overlay of C-lobes of the structure complex of MAPKAPK2-ΔAiα with Bromo-MK-25 in grey and the inactive MAPKAPK2 (PDB 7NRY, Croll et al., 2021) represented in magenta cartoon. Secondary structure nomenclature is taken from Meng et al., 2002.

Together, these rearrangements generate a previously unobserved cavity. The location and shape of this pocket explain the state-dependence of MK-25 ligand binding: in the autoinhibited conformation, the site is either simply occluded by the autoinhibitory helix or not fully formed, because the surrounding C-lobe elements have not adopted the binding-competent arrangement.

## 4. Discussion

This study defines the structural basis for recognition of an allosteric furan-2-carboxamide inhibitor by activated MAPKAPK2. Although MAPKAPK2 has been studied extensively as a therapeutic target (Gaestel et al., 2006; Fiore et al., 2016), prior structural work has largely focused on ATP-competitive ligands (Underwood et al., 2003; Anderson et al., 2009a and 2009b; Fiore et al., 2016). The present structure therefore fills an important gap by revealing a non-canonical ligand-binding pocket in the MAPKAPK2 C-lobe and by linking formation of this pocket to activation-associated conformational rearrangements.

A central finding is that binding of MK-25 is strongly state-dependent. The ligand shows no detectable interaction with the inactive non-phosphorylated MAPKAPK2, whereas p38α-activated or constitutively active constructs bind the ligand with submicromolar affinity. These observations support a mechanism in which autoinhibition prevents formation or exposure of the allosteric site. Relief of autoinhibition, whether through phosphorylation-dependent activation or – as done in this work – through truncation of the autoinhibitory helices, permits reorganization of the αD and αG helices and renders the allosteric site accessible. This state-dependence is noteworthy from an inhibition mechanism point of view, as it indicates that the allosteric site is formed only in a conformation less broadly conserved across the kinome than the ATP-binding pocket.

The location of Bromo-MK-25 in the MAPKAPK2 C-lobe bears a notable resemblance to the binding site of the GluN2B peptide substrate in activated CaMKII kinase (PDB: 7UJT) (Ozden et al., 2022). Although both kinases have different activation mechanisms (Rellos et al., 2010), they are both members of the CaMK superfamily and share a similar substrate recognition specificity (Stokoe et al., 1993), as they both phosphorylate several common protein substrates (i.e. both phosphorylate tyrosine hydroxylase (Campbell et al., 1986) and glycogen synthase (Woodgett et al., 1982)). Structural superposition shows that the MK-25 analogue occupies a region structurally similar to that engaged by the GluN2B substrate in CaMKII, with the piperazine moiety of Bromo-MK-25 positioned analogously to the Arg1295 residue of the GluN2B peptide (Figure 5). Moreover, the bromo-phenyl group occupies the same pocket as Leu1298 of the substrate GluN2B. These observations support a structural overlap between the Bromo-MK-25 binding site and the substrate-recognition surface of MAPKAPK2. Notably, enzymatic analysis of a related compound from this series showed non-competitive inhibition with respect to the peptide substrate (Huang et al., 2011), indicating that inhibition does not arise from simple steric exclusion of substrate binding. This suggests a more nuanced allosteric mechanism, in which occupancy of this pocket may impair catalytic efficiency or substrate positioning without fully preventing substrate engagement.

**Figure 5.**
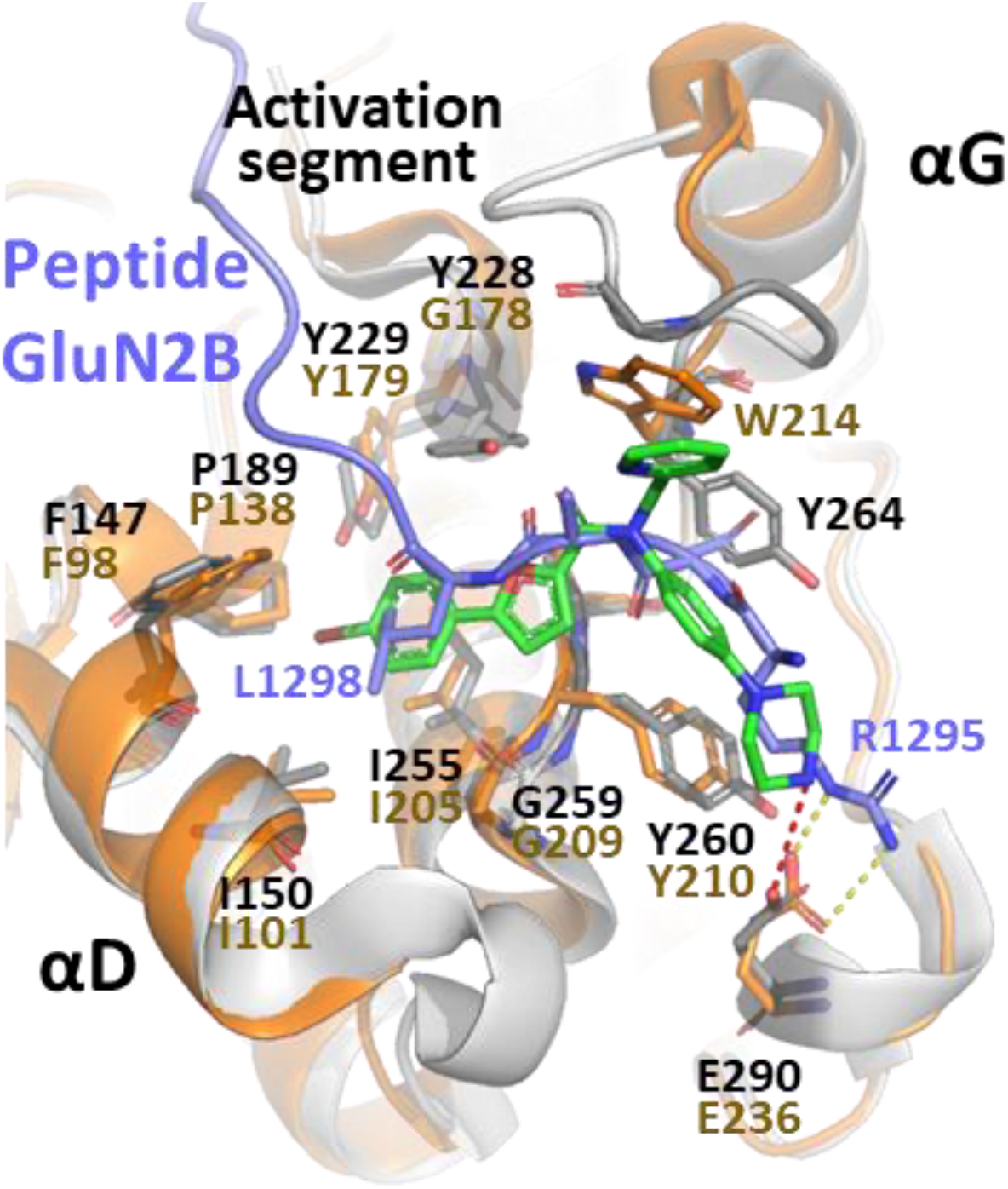
Overlay of Bromo-MK-25 and GluN2B binding sites in MAPKAPK2 and CaMKII. MAPKAPK2 and CaMKII (PDB ID 7UJT) are represented in grey and brown cartoon respectively. The key MAPKAPK2 binding site residues are shown in grey sticks with label in black. The related residues for CaMKII, with labels, are in brown. Bromo-MK-25 is represented in green and the peptide GluN2B in blue. Hydrogen bonds from piperazine in dash red and from Arg1295 in dash yellow.

The crystal structure complex is consistent with previously published SAR observations for the furan-2-carboxamide series (Huang et al., 2011; Huang et al., 2012). In this respect, the crystallographic data provide direct support for assigning this site as the furan-2-carboxamide binding location. The main advance of the present work is therefore the identification and structural characterization of a ligandable allosteric pocket in activated MK2.

Two lines of evidence support the physiological relevance of the observed conformational rearrangement, while also indicating its limits. Firstly, the reorganization of the MAPKAPK2 C-lobe is compatible with regulatory behaviour described for related kinase family members, including CaMKII (Rellos et al. 2010) and RSK2 (Malakhova et al., 2008), suggesting that the structural transitions captured here are mechanistically meaningful rather than a crystallographic artefact. Secondly, the allosteric binding mode was defined only in the presence of an ATP-site ligand (PF-3644022) and an engineered constitutively active construct, so the extent to which the observed pocket geometry matches that of fully phosphorylated wild-type MAPKAPK2 in solution remains to be established. The ITC data argue against the interaction being a crystallisation artefact, but future structural studies will be needed to clarify the mechanistic coupling between orthosteric occupancy, activation state, and allosteric-site formation.

Overall, the data identifies an active-state C-lobe pocket that is ligandable, structurally distinct from the ATP site, and compatible with previously reported SAR for the MK-25 series. These findings establish a structural framework for understanding allosteric recognition in MAPKAPK2 and expand the known conformational repertoire of this therapeutically relevant kinase.

## 5. Associated content

### Supporting information

Details of the protein sequences; The fitted and omit map for Bromo-MK25 bound in MAPKAPK2:PF-3644022:Bromo-MK-25 (Figure S1); refinement of statistics for X-ray crystal structures (Table T1); supplementary tables and figures.

### Accession code

The X-ray crystal structure was deposited into the Protein Data Bank with the PDB code 6T8X.

## CRediT authorship contribution statement

**Edward Beaumont:** Coordination of the research, Supervision, Investigation, Methodology, Formal analysis, Data curation, Visualization, Writing – original draft. **Stephanie Duclos:** Formal analysis and manuscript revision. **Sandrine Thieffine:** Investigation, Methodology. **Rahul Yadav:** Investigation, Methodology. **John Barker:** Conceptualization, Supervision, Project administration and manuscript revision.

## Notes

All authors are currently or have been employees of Evotec (UK) Ltd at the time of the study.

## SUPPLEMENTAL INFORMATION

**Figure S1.**
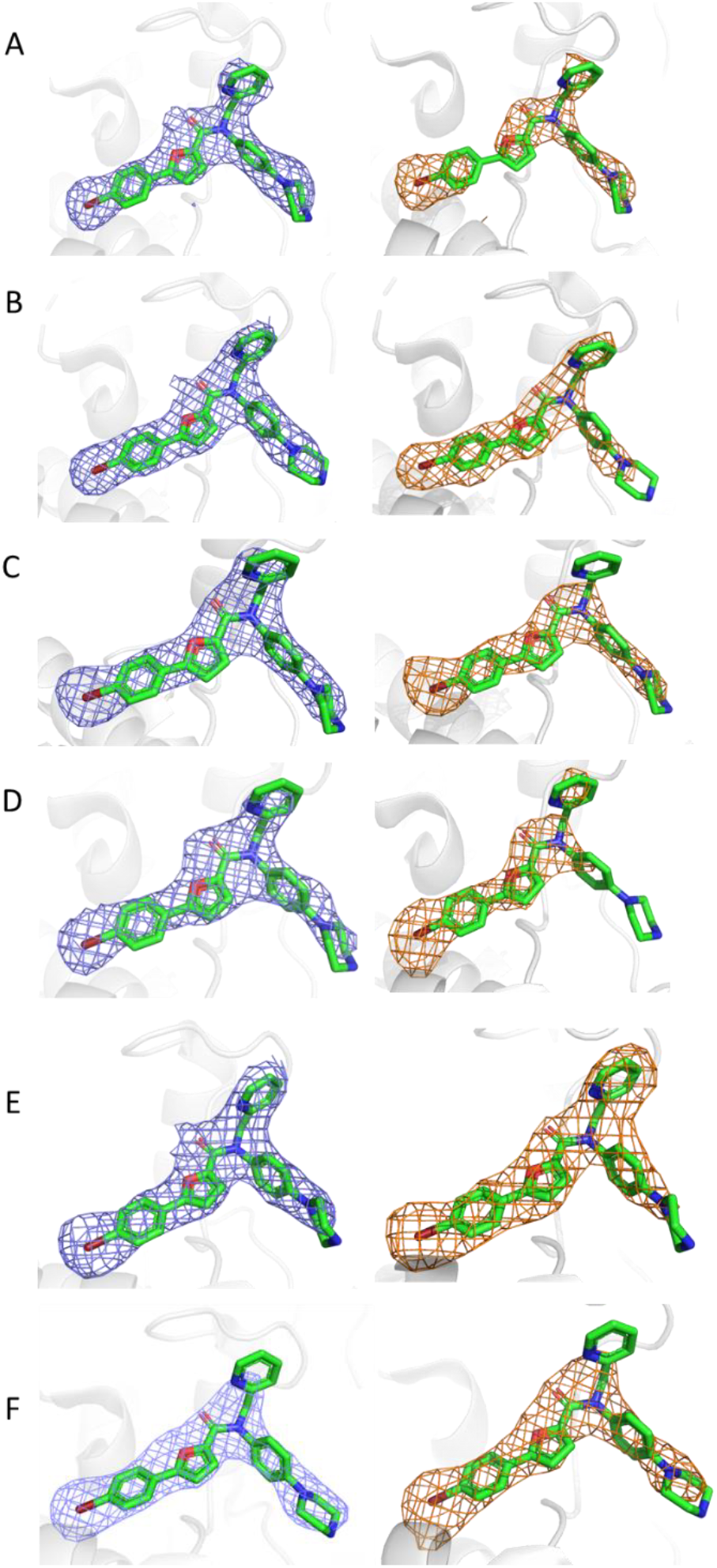
The fitted and omit map for Bromo-MK-25 bound in MAPKAPK2:PF-3644022:Bromo-MK-25. The 2Fo-Fc map calculated (blue, contoured at 1 σ) after final refinement, with model including the ligands for chain A, B, C, D, E and F. The Fo-Fc map calculated (orange, contoured at 2.5 σ) prior to the addition of ligands to the model for chain A, B, C, D, E and F.

**Table T1 - Crystallographic table**

**Table 1.**
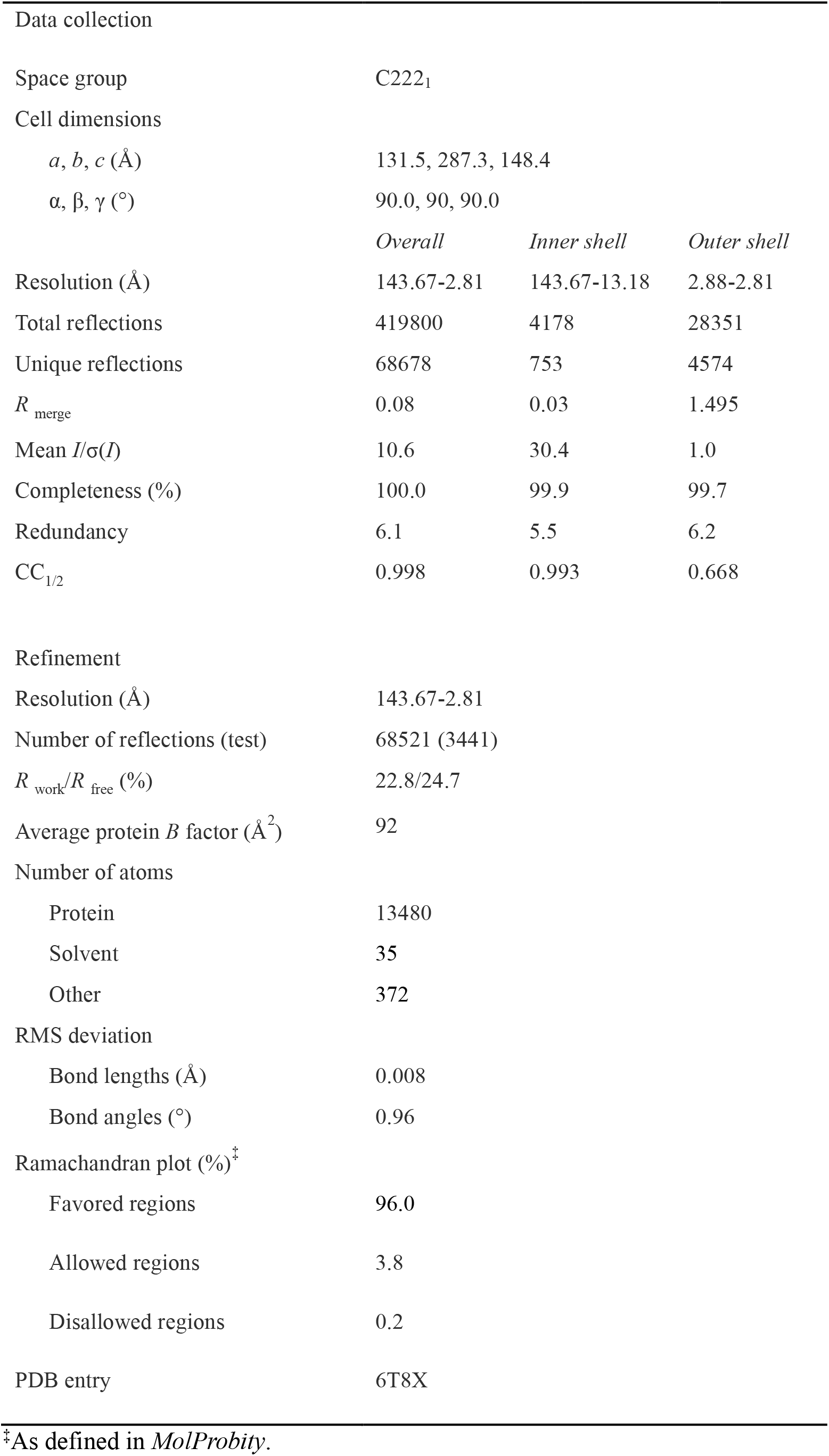
Data-collection and refinement statistics of MAPKAPK2:PF-3644022:Bromo-MK-25 complex.

### Crystal structure of activated MAPKAPK2 kinase reveals a C-lobe allosteric pocket for furan-2-carboxiamide inhibitors

Edward Beaumont, Stephanie Duclos, Sandrine Thieffine, Rahul Yadav, and John Barker

### Protein sequences

#### Crystallographic construct: MAPKAPK2 Q41-T338 T222E T334E

MGSSHHHHHHSSGENLYFQGSQQFPQFHVKSGLQIKKNAIIDDYKVTSQVLGLGINGKVLQIFNKRTQEKFALKMLQDCPKARREVELHWRASQCPHIVRIVDVYENLYAGRKCLLIVMECLDGGELFSRIQDRGDQAFTEREASEIMKSIGEAIQYLHSINIAHRDVKPENLLYTSKRPNAILKLTDFGFAKETTSHNSLTEPCYTPYYVAPEVLGPEKYDKSCDMWSLGVIMYILLCGYPPFYSNHGLAISPGMKTRIRMGQYEFPNPEWSEVSEEVKMLIRNLLKTEPTQRMTITEFMNHPWIMQSTKVPQ EPLHT

### Constructs used for the ITC experiments

#### Construct MAPKAPK2 H47-H400

MGSSHHHHHHSSGENLYFQGSHVKSGLQIKKNAIIDDYKVTSQVLGLGINGKVLQIFNKRTQEKFALKMLQDCPKARREVELHWRASQCPHIVRIVDVYENLYAGRKCLLIVMECLDGGELFSRIQDRGDQAFTEREASEIMKSIGEAIQYLHSINIAHRDVKPENLLYTSKRPNAILKLTDFGFAKETTSHNSLTTPCYTPYYVAPEVLGPEKYDKSCDMWSLGVIMYILLCGYPPFYSNHGLAISPGMKTRIRMGQYEFPNPEWSEVSEEVKMLIRNLLKTEPTQRMTITEFMNHPWIMQSTKVPQTPLHTSRVLKEDKERWEDVKEEMTSALATMRVDYEQIKIKKIEDASNPLLLKRRKKARALEAAALAH

#### Construct MAPKAPK2 M1-T338

MGSSHHHHHHSSGENLYFQGSMLSNSQGQSPPVPFPAPAPPPQPPTPALPHPPAQPPPPPPQQFPQFHVKSGLQIKKNAIIDDYKVTSQVLGLGINGKVLQIFNKRTQEKFALKMLQDCPKARREVELHWRASQCPHIVRIVDVYENLYAGRKCLLIVMECLDGGELFSRIQDRGDQAFTEREASEIMKSIGEAIQYLHSINIAHRDVKPENLLYTSKRPNAILKLTDFGFAKETTSHNSLTTPCYTPYYVAPEVLGPEKYDKSCDMWSLGVIMYILLCGYPPFYSNHGLAISPGMKTRIRMGQYEFPNPEWSEVSEEVKMLIRNLLKTEPTQRMTITEFMNHPWIMQSTKVPQTPLHT

**Constructs used for phosphorylation of kinases**; Constitutively active MKK6-DD was used to phosphorylate p38α. Then, phosphorylated p38α was used to phosphorylate MAPKAPK2.

#### Construct MKK6-DD M1-D334 S207D T211D

MGSSHHHHHHSSGENLYFQGSMSQSKGKKRNPGLKIPKEAFEQPQTSSTPPRDLDSKACISIGNQNFEVKADDLEPIMELGRGAYGVVEKMRHVPSGQIMAVKRIRATVNSQEQKRLLMDLDISMRTVDCPFTVTFYGALFREGDVWICMELMDTSLDKFYKQVIDKGQTIPEDILGKIAVSIVKALEHLHSKLSVIHRDVKPSNVLINALGQVKMCDFGISGYLVDDVAKDIDAGCKPYMAPERINPELNQKGYSVKSDIWSLGITMIELAILRFPYDSWGTPFQQLKQVVEEPSPQLPADKFSAEFVDFTSQCLKKNSKERPTYPELMQHPFFTLHESKGTDVASFVKLILGD

#### Construct murine p38α S2-S360 based on White *et al*., 2007

MAHHHHHHSQERPTFYRQELNKTIWEVPERYQNLSPVGSGAYGSVCAAFDTKTGHRVAVKKLSRPFQSIIHAKRTYRELRLLKHMKHENVIGLLDVFTPARSLEEFNDVYLVTHLMGADLNNIVKCQKLTDDHVQFLIYQILRGLKYIHSADIIHRDLKPSNLAVNEDCELKILDFGLARHTDDEMTGYVATRWYRAPEIMLNWMHYNQTVDIWSVGCIMAELLTGRTLFPGTDHIDQLKLILRLVGTPGAELLKKISSESARNYIQSLAQMPKMNFANVFIGANPLAVDLLEKMLVLDSDKRITAAQALAH AYFAQYHDPDDEPVADPYDQSFESRDLLIDEWKSLTYDEVISFVPPPLDQEEMES

